# Conservation of declining Florida ecosystems is critical for an at-risk butterfly

**DOI:** 10.64898/2026.08.04.742755

**Authors:** Ava Johnson, Rachel L. Walsh, Bert Foquet, Jaret C. Daniels, Robert P. Guralnick, Akito Y. Kawahara

**Affiliations:** McGuire Center for Lepidoptera and Biodiversity, Florida Museum of Natural History, University of Florida, Gainesville, FL 32611, USA; Department of Geography, University of Florida, Gainesville, FL 32611, USA; School of Natural Resources and Environment, Institute of Food and Agricultural Sciences, College of Agricultural and Life Sciences, University of Florida, Gainesville, FL 32611, USA; Division of Informatics, Florida Museum of Natural History, University of Florida, Gainesville, FL 32611, USA

**Keywords:** habitat, Hesperiidae, land use, Lepidoptera, protected areas, spatial ecology

## Abstract

Assessing the availability of protected habitats for at-risk species is needed to determine how the spatial distribution of land cover and land use shapes conservation outcomes. The Loammi skipper (*Atrytonopsis loammi*) is a nonmigratory, prairie-associated butterfly that has experienced a significant reduction in its formerly widespread southeastern United States distribution over the past several decades. Observations in recent years have been limited to a small number of isolated Florida populations, and it is unclear how much suitable habitat remains and what proportion of that habitat is protected. Here we used publicly available community science data and collected specimens to identify the predominant land cover types occupied by *A. loammi*. We then quantified the extent to which each of these land cover types overlap with protected areas to estimate the proportion of the butterfly’s current distribution that occurs on managed conservation lands. Our findings show that *A. loammi* relies heavily on protected lands, with over half of the total area of the most strongly associated habitats located within protected areas. Given the continued expansion of human-modified landscapes in Florida and the reliance of *A. loammi* on protected lands, the long-term persistence of *A. loammi* will likely depend on the conservation and effective management of its remaining suitable habitat.

**Implications for insect conservation:** Our results highlight the critical role of protected areas in sustaining at-risk insect populations, particularly for species with limited dispersal and shrinking ranges. More broadly, they suggest that conserving and restoring protected habitats while maintaining connectivity are essential strategies for effective insect conservation in rapidly developing regions.

## Introduction

Insect populations are declining worldwide (Cardoso et al. 2020; Dirzo et al. 2014; Forister et al. 2021; Wagner et al. 2021). A recent global meta-analysis by Wagner et al. (2021) found that the rate of decline is about 1-2% annually. This is a significant concern because insects provide many essential ecosystem services, including pollination, waste decomposition, and nutrient cycling; form the foundation of ecological networks across nearly every terrestrial ecosystem; and support economic sectors ranging from agriculture to biotechnology (Dicke 2018; Jankielsohn 2018; Losey & Vaughan 2006; Scudder 2017; Verma et al. 2023). Among the many factors contributing to insect declines, land use changes (e.g., natural habitat conversion to urban or agricultural land) and habitat loss and degradation are especially notable (Forister et al. 2019; Maes & Van Dyck 2001; Sánchez-Bayo & Wyckhuys 2019; Wagner et al. 2021). Therefore, investigating how land cover and land use coincide with the geographic distribution of at-risk insects is critical to better understand species’ vulnerability.

Conservation areas are known for their importance in protecting vulnerable populations (Nowakowski et al. 2023; Vellak et al. 2009). While this has been extensively studied in plants and vertebrates, insects and other invertebrates generally receive less consideration when establishing protected areas, and their dependence on conservation lands is less understood (Chowdhury et al. 2023a; Chowdhury et al. 2023b; Sinclair et al. 2025). For example, open-canopy habitats tend to hold higher insect biomass and diversity but are less likely to be protected than forested areas (Sinclair et al. 2025). Chowdhury et al. (2023a) estimated that about 76% of insects fail to meet the minimum target coverage of protected areas needed for adequate long-term protection, which are set according to species’ geographic ranges. Despite receiving less consideration in the establishment of protected areas, studies show that insects benefit from their conservation. Gillingham et al. (2015) found that about 80% of historically resident butterfly and odonate species in the UK were more abundant within protected areas, and about 61% were more abundant within protected areas recently colonized due to climate tracking. These studies highlight the importance of understanding critical habitat areas for insects and ensuring that insects are included when designating protected areas.

Florida is the third most populous U.S. state and the second fastest growing in terms of absolute population (U.S. Census Bureau 2026a; U.S. Census Bureau 2026b). Land cover changes have occurred rapidly in Florida over the past century as a result of conversion of natural areas to agriculture and pasturelands, urbanization, mining, wetland drainage, fire suppression, climate change, and exotic species invasions (Volk et al. 2017). Dry prairie has faced the greatest decline in percent area, with a 25% reduction between the late 1980s and 2003 (Kautz 2007). Meanwhile, pinelands experienced the greatest decline in total area, losing 0.24 million ha during the same period (Kautz 2007). Both dry prairies and pinelands are critical habitats for many at-risk Florida species (Florida FWC 2019), making their declines especially concerning. These pressures are expected to intensify as Florida’s population is projected to increase by 10 million residents by 2070, driving continued urban expansion and land use change (Carr & Zwick, 2016). Consequently, Florida represents an important case study for understanding how habitat loss and land use changes impact at-risk insects.

Butterflies, one of the most familiar and well-liked insect groups among the general public, have shown declines across the globe (Edwards et al. 2025; Maes & Van Dyck 2001; Warren et al. 2021). In Europe, monitoring efforts for 16 countries have shown an overall 39% decline of grassland butterflies since 1990 (Warren et al. 2021). Similarly, U.S. butterfly abundance declined by around 1.3% annually between 2000 and 2020 (Edwards et al. 2025). Because they are charismatic and well-studied, butterflies are an excellent ambassador group for showcasing the importance of action to address insect declines (Kawahara et al. 2021). Investigating and raising awareness about the ecological needs of local species can promote conservation of critical habitat for butterflies along with a variety of at-risk insects.

The Loammi skipper (*Atrytonopsis loammi*) is a fairly large, nonmigratory grass skipper (Hesperiinae), ranked globally imperiled by NatureServe (NatureServe 2026) and listed as a Species of Greatest Conservation Need by the Florida Fish and Wildlife Conservation Commission (Florida FWC 2019; Schweitzer et al. 2018). An approximately 90% decline in the species’ geographic range over the past century has been largely attributed to habitat loss (NatureServe 2026). Historically found throughout the southeastern United States, the species is currently only known from a small number of isolated Florida populations (Schweitzer et al. 2018). The suitable habitat has been described as open-canopy pine savannas, high pine, scrub, dry prairies, pine flatwoods, and freshwater non-forested wetlands with high densities of the known host plant, lopsided Indiangrass (*Sorghastrum secundum*) (Florida FWC 2019; Schweitzer et al. 2018). The majority of associated habitats are fire-maintained (Florida FWC 2019; Schweitzer et al. 2018) and habitat degradation through fire suppression and alterations to fire regime (e.g., scale, intensity, and return interval) is considered a significant threat. Therefore, active management is needed to maintain natural fire regimes (Adamidis et al. 2019; Florida FWC 2019; Jue et al. 2022; Schweitzer et al. 2018). It is unclear how much suitable habitat for the Loammi skipper remains, or how much of the suitable habitat occurs within managed conservation lands. Here we address the following questions relevant to the conservation of this species: (1) which land cover classes are most commonly associated with the Loammi skipper, and (2) what proportion of these land cover classes is located within Florida conservation lands?

## Materials and Methods

### Occurrence data

Occurrence records for *A. loammi* were gathered from two datasets: (1) GPS locations were recorded for genetic samples opportunistically collected from nine known occurrence sites across the species’ current range from March-September 2023 (n=155), and (2) all validated and unobscured iNaturalist observations from community scientists who joined an iNaturalist project were downloaded on June 10th, 2026 (n=455). For the first dataset, experts were consulted in an intentional effort to sample from all known occupied sites. Both datasets were imported into R v 4.5.2 (R Core Team 2025) and merged together. To maintain temporal consistency with land cover data, the resulting dataset was filtered by date of observation to retain only records within 10 years of 2025. Occurrence records were then filtered to retain only records with ≤ 75 m coordinate uncertainty. The 75-m threshold was selected to keep coordinate uncertainty within a reasonable estimate of dispersal distance. The resulting coordinate uncertainty for iNaturalist data was 1-72 m with a mean of 26 m. For collections data, the GPS locations were recorded on an iPhone 11 in open-canopy environments with clear weather, which yields coordinate accuracy levels comparable with recreation-grade GPS receivers (7-13 m) (Merry & Bettinger 2019). The data were then spatially thinned by 150 m using *spThin* v 0.2.0 to reduce spatial autocorrelation (Aiello-Lammens et al. 2015) (Table S1, Online Resource 1). A 1-km spatial thin was also performed which resulted in largely identical habitat rankings compared to the 150-m thin but retained only 50 occurrences (Table S2, Online Resource 1). These methods produced a final dataset with 117 occurrences across the skipper’s known range (Collections: 57; iNaturalist: 60).

### Land cover data

Land cover data were imported from the Florida Cooperative Land Cover (CLC) raster (v 4.0, November 2025) (Florida FWC & FNAI 2025). Produced as a collaboration between the Florida Fish and Wildlife Conservation Commission (Florida FWC) and the Florida Natural Areas Inventory (FNAI) by combining ground-truth mapping and expert review of aerial imagery and existing data sources, this resource represents the most comprehensive and up-to-date land cover mapping dataset for Florida (Florida FWC & FNAI 2025). The dataset uses the Florida Land Cover Classification System, which is composed of a hierarchy of increasingly specific land cover classes modeled after the Florida Land Use, Cover, and Forms Classification System (Florida DOT 1999). In this study, we used the more generalized STATE level raster, consisting of 75 land cover classes within Florida.

### Background sampling

We accounted for sampling biases by first estimating a region where *A. loammi* could occur and then using surrogate sampling of related taxa within this region to quantify bias. Using R v 4.5.2 (R Core Team 2025), we delimited an accessible area (i.e., an estimate of the total area that *A. loammi* can access), using dispersal distances of closely related skippers to estimate dispersal ability (Leidner & Haddad 2011; U.S. Fish and Wildlife Service 2021). We estimated dispersal at 0.4 km per 3 days and extrapolated this rate over 10 years, assuming two broods per year and a four-week adult lifespan. We used the resulting estimate (75 km) with an added 5-km buffer to create an alpha hull around the pre-thinned occurrence records with *rangeBuilder* v 2.2 (Davis Rabosky et al. 2016) (Figure S1, Online Resource 1). We generated a bias layer consisting of a surrogate measure of sampling intensity by downloading all validated, unobscured butterfly (Lepidoptera: Papilionoidea) occurrences from iNaturalist within the accessible area recorded within 10 years of 2025 with ≤ 75 m coordinate accuracy (n = 42,259). We operationalized our bias layer by using *terra* v 1.8.60, to sample 10,000 background points across the accessible area weighted by the bias layer, which resulted in a control that accounts for observer effort (Hijmans et al. 2020; Phillips et al. 2009) (Figure S1, Online Resource 1).

### Habitat rankings

To assess which land cover classes are most commonly associated with *A. loammi*, we produced habitat rankings in Python v 3.12.10 using the libraries *NumPy* v 2.4.0 (Harris et al. 2020), *pandas* v 3.0.1 (The pandas development team 2024; McKinney 2010), *GeoPandas* v 1.1.2 (Jordahl et al. 2020), *Rasterio* v 1.5.0 (Gillies 2013), and *Shapely* v2.1.2 (Gillies et al. 2025). A 75-m buffer that corresponded with GPS uncertainty was generated around each occurrence. This method was used to account for the fact that the true location might fall anywhere within the buffer and the assumption that *A. loammi* can disperse to anywhere within this area. Land cover classes within each buffer were extracted from the Florida CLC map, and their respective proportions were calculated based on pixel counts. We also established a 75-m buffer around each of the background points and extracted land cover using the same method.

A habitat index value was calculated for each land cover class by dividing the proportion of pixels for a given land cover class within all 75-m buffers for *A. loammi* occurrence records by the proportion of pixels for that class within all 75-m buffers for the background points (Davis et al. 1994). The aim of creating this index was to standardize the frequency of occurrence within each land cover class by butterfly observer effort and exposure to each land cover class, revealing a clearer picture of habitat association. Land cover classes were then ranked based on the habitat index to identify the top 10 habitat associations of *A. loammi*. Using this method, 75-m uncertainty buffers around occurrence records likely included both suitable habitat types as well as those that are less suitable but adjacent to suitable habitat types. However, by aggregating the total proportion of each land cover class across all 117 *A. loammi* occurrences, we believe that we are using the best available information to capture a signal of habitat association.

### Conservation lands data

The Florida Land Managed Areas shapefile was downloaded from FNAI (FNAI 2026). This database consists of over 3,000 managed areas, including national parks, state forests, wildlife management areas, and local and private preserves. These areas are characterized by having maintained most of their natural attributes, being significantly undeveloped, and having a management agency formally committed to conserving their natural condition (FNAI 2026).

### Proportion of habitats protected

The proportion of each habitat protected was calculated by overlaying the shapefile of Florida Land Managed Areas with the CLC raster and counting the raster pixels for each land cover class whose centers fell within conservation boundaries. Data were processed using Python v 3.12.10 with the libraries *NumPy* (Harris et al. 2020), *pandas* (The pandas development team 2024; McKinney 2010), *Rasterio* (Gillies 2013), and *SciPy* v1.17.0 (Virtanen et al. 2020).

## Results

### Habitat associations

Our occurrence datasets included records from multiple geographically separate *A. loammi* populations (Figure 1): a Florida panhandle population located in Apalachicola National Forest (Figure 1a), a central Florida metapopulation spanning several conservation areas (Figure 1b), and a south Florida population in Jonathan Dickinson State Park (Figure 1c). Based on our index, the most common habitat association was Dry Prairie followed by Palmetto Prairie (Figure 2a; Table S1, Online Resource 1). Mesic Flatwoods, Cultural - Riverine, and Scrubby Flatwoods were the next most common habitats, although each received less than half the index score of Dry Prairie (Figure 2a; Table S1, Online Resource 1). Over half of *A. loammi* occurrences in this study (64 out of 117) were predominately associated with Mesic Flatwoods (the majority land cover class within the 75-m buffer area was Mesic Flatwoods). However, pine flatwoods are the most common habitat in Florida (Florida FWC 2019), and thus, higher sampling frequency likely reduced the index score of the corresponding land cover classes as a result of standardization. Finally, the list of top 10 habitats also included Cultural - Terrestrial, Dome Swamp, Prairies and Bogs, Isolated Freshwater Marsh, and Scrub (Figure 2a; Table S1, Online Resource 1). Land cover class descriptions can be found in Kawula et al. (2025).

**Fig. 1.**
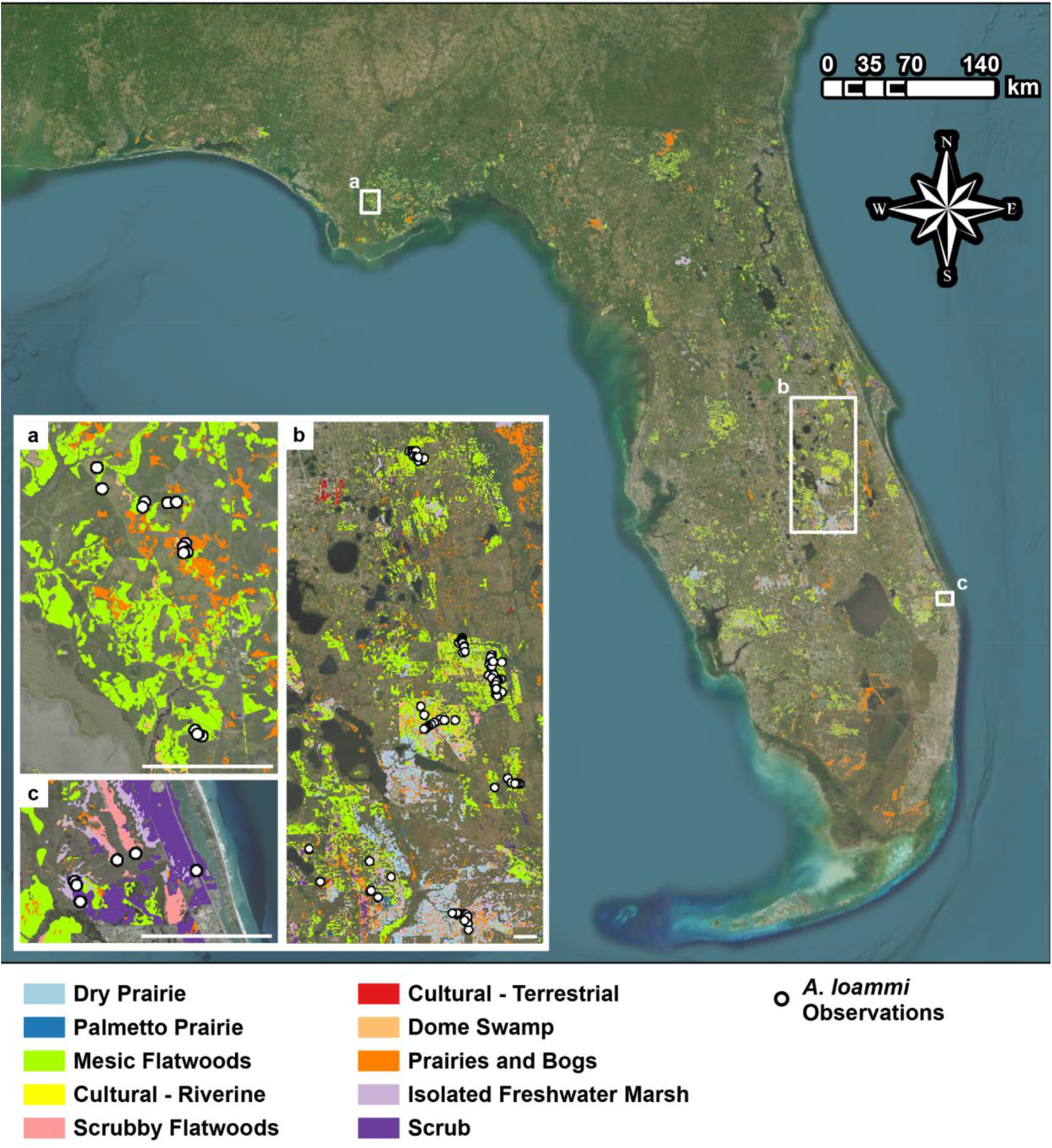
Map of Florida showing the top 10 habitat associations of *A. loammi* as ranked by habitat index. Insets represent the three geographically separate populations of *A. loammi*: (a) Florida panhandle, (b) central Florida, (c) south Florida. The scale bar in the lower right corner of each inset is 5 km. Maps created in ArcGIS Pro with layout completed in Adobe Illustrator. Basemap source: Esri, Vantor, Earthstar Geographics, and the GIS User Community

**Fig. 2.**
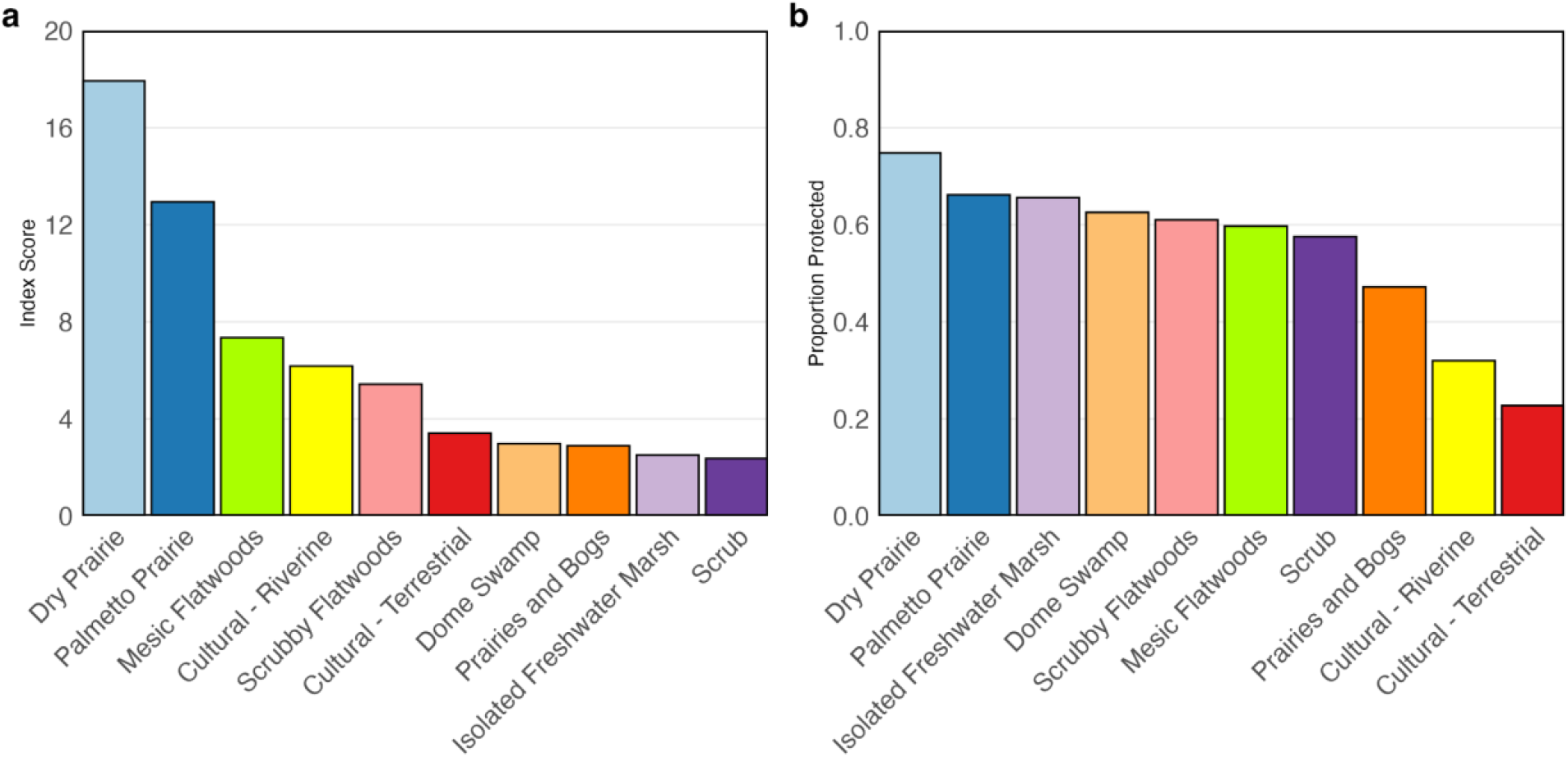
Index-based habitat association rankings for *A. loammi* (a) and habitat proportions within protected areas (b). Colors associated with each habitat type match the colors used in Figure 1. Figure created using ggplot2 in R v 4.5.2

### Reliance on protected lands

In total, 56% of the total area covered by the top 10 habitats within which *A. loammi* has been recorded fell within Florida conservation lands (Figure 2b; Table S1, Online Resource 1). All *A. loammi* records included in our study were observed within protected areas (Figure S2, Online Resource 1). The total area in Florida of the top 10 habitats as of November 2025 is 3.75 million acres (Figure 1; Table S1, Online Resource 1).

Dry Prairie, the highest ranked habitat in our index (Figure 2a), also ranked highest in terms of proportion protected (Figure 2b). There are 152,809 total acres of Dry Prairie in Florida (Figure 1; Table S1, Online Resource 1) and nearly 75% of this area (114,607 acres) fell within Florida Land Managed Areas (Figure 2b; Table S1, Figure S2, Online Resource 1). Palmetto Prairie, which was second in our habitat ranking and second in terms of proportion protected (Figure 2), had the smallest total area within Florida of the top 10 habitats (Figure 1; Table S1, Online Resource 1). There are 18,798 total acres attributed to this land cover class and 66% (12,406 acres) are protected (Figure 1, 2b; Table S1, Figure S2, Online Resource 1). Mesic Flatwoods ranked third in our habitat index and sixth in terms of proportion protected (Figure 2), with over 1.5 million total acres and 897,955 acres protected (59.7%) (Figure 2b; Table S1, Figure S2, Online Resource 1). The two human-modified land use types (Cultural - Riverine, Cultural - Terrestrial) that appeared in our top 10 habitats (Figure 2a) had the lowest proportions protected when compared with the top 8 natural habitats (Figure 2b; Table S1, Figure S2, Online Resource 1). Prairies and Bogs was the only top 8 natural habitat under 50% protected (Figure 2b; Table S1, Figure S2, Online Resource 1).

## Discussion

This study is the first to systematically evaluate habitat associations for *A. loammi* using occurrence data from across the species’ current known range. We confirmed previously published natural history descriptions (Florida FWC 2019; Schweitzer et al. 2018), revealing Dry Prairie and Palmetto Prairie as the top-ranked habitats, with Mesic Flatwoods and Scrubby Flatwoods also shown as highly ranked habitat associations. Mesic Flatwoods was predominately associated with the greatest number of *A. loammi* observations in our study (n = 64). However, its index score was lowered due to standardization based on sampling effort, elevating Dry Prairie and Palmetto Prairie as the top habitats when accounting for effort.

This study also demonstrated the importance of protected lands for *A. loammi*. All *A. loammi* observations, as well as 56% of the total area encompassed by the 10 highest ranked habitat types, occurred within Florida Land Managed Areas, including federal, state, local, and private preserves (Figure 2; Table S1, Figure S2, Online Resource 1). Dry Prairie, the top-ranked habitat in our analysis, showed an even stronger association with protected lands, with approximately 75% of its total area occurring within protected areas across Florida (Figure 2). Our collections represented an intentional effort to sample all known locations of *A. loammi*, which all occur within conservation lands; therefore, there may be a slight bias in our dataset towards conservation lands. Community science observers might similarly show a bias towards searching for butterflies within conservation areas, but they may likewise be biased towards areas closer to urban and residential areas where people live, which are outside conservation lands. To account for these potential biases, we standardized our habitat index using a measure of sampling effort for all butterfly species, which reduces the influence of any oversampled areas (e.g., conservation-associated land cover classes) in our habitat index.

Although the high proportion of *A. loammi* habitat within protected areas may help safeguard the species’ future, this pattern may also reflect habitat loss outside protected lands due to the rapid expansion of human-modified landscapes in Florida. Continued habitat loss would not only reduce the total area of suitable habitat but also reduce connectivity between already fragmented habitat patches (Haddad et al. 2015; Peled et al. 2026). Protecting and actively managing top habitats while maintaining connectivity is likely critical for the long-term survival of this species.

Two human-modified land use types appeared in the top 10 habitats (Cultural - Riverine, Cultural - Terrestrial). Cultural land cover types are defined by substantial modification of substrate, streamflow, morphometry, or biological composition due to human activities (Kawula et al. 2025). As above, it is important to note that occurrence records used in this study were opportunistically collected, making them susceptible to bias. Although we attempted to control for bias, it is possible that our data may contain biases toward areas that are easily accessible to observers, which might result in human-modified land uses ranking higher in our index.

Additionally, our data show that all *A. loammi* records either fell within or in close proximity to top-ranked natural habitats. In total, seven *A. loammi* observations were classified as > 75 m outside the top 8 natural habitats, and estimated distances were on average about 212 m away from the nearest top 8 natural habitat. Although *A. loammi* dispersal ability has not been directly investigated, a recent study reported a maximum dispersal distance of 400 m over several days for the Crystal skipper (*Atrytonopsis quinteri*), a similarly sized and closely related species (Leidner & Haddad 2011). The maximum distance estimated between an *A. loammi* occurrence and a top-ranked natural habitat was 344 m (Table S3, Online Resource 1), which is within the reported dispersal range for *A. quinteri*. Therefore, *A. loammi* presence within human-modified and lower ranked areas could be due to dispersal rather than persistence.

Although our data suggest that the skippers can disperse between human-modified areas and nearby highly ranked natural habitats, it is possible that *A. loammi* uses the human-modified land use areas. Disturbed areas often contain large populations of ruderal flowering plants, such as yellow thistle (*Cirsium horridulum*), which is commonly used as a nectar source by *A. loammi* (Schweitzer et al. 2018). The Cultural - Terrestrial land cover class includes Mowed Grass, Vegetative Berm, and Highway Rights of Way, which are all characterized by early successional, herbaceous vegetation (Kawula et al. 2025). During field surveys that produced data for this study, *A. loammi* was frequently observed using open-canopy roadsides for nectar, which has been documented for many insect species (Kimmel et al. 2024; Wagner et al. 2014). One possibility is that *A. loammi* individuals move between nectar-rich and host-rich areas in pursuit of different resources. Availability of both larval host plants and nearby nectar sources is likely important for *A. loammi* persistence. Management of human-modified landscapes to maintain herbaceous plant communities can also benefit at-risk insects (Kimmel et al. 2024).

This study uses community science and collected specimen records that are opportunistic in nature and may include repeated sampling of known populations and heavier sampling in areas with ease of access to human observers; however, we accounted for these biases using spatial point thinning and standardization using a bias layer that accounted for sampling effort. Nonetheless, future studies using systematically collected occurrence data could provide a more complete picture of *A. loammi* distribution and habitat requirements. Additionally, future studies could empirically quantify *A. loammi* dispersal ability, which was not directly measured in this study. As this species inhabits fire-maintained ecosystems, more research is needed to understand which fire regimes are most beneficial (e.g., burn unit sizes, seasonality, return interval). Ecological niche modeling studies would also be beneficial to further characterize suitable habitat. In general, more data on *A. loammi* ecology, population dynamics, and life history are needed to inform management of this at-risk species.

Our study highlights the value of community science. The majority of *A. loammi* occurrence records used in this study were obtained from iNaturalist, a software application that allows users to record natural history observations in a publicly accessible database (iNaturalist 2026). Participation with community science platforms such as iNaturalist has dramatically increased the amount of natural history data available to researchers. This has not only made studies on under-researched species such as *A. loammi* more feasible but also allows community members to make meaningful contributions to scientific research (Mason et al. 2025).

In conclusion, this is the first study to directly quantify habitat associations for the at-risk Loammi skipper, confirming previously published natural history accounts that described dry prairie and pine flatwoods as critical habitats (Florida FWC 2019; Schweitzer et al. 2018). Additionally, our findings suggest *A. loammi* may disperse between highly ranked natural habitats and nearby human-modified landscapes, which it may utilize for nectar resources. A major finding of this study is the high proportion (56%) of top-ranked *A. loammi* habitats located within Florida protected areas, which indicates that this species is highly dependent on conservation lands. Although the high percentage of protected habitat is encouraging, this may also indicate that suitable habitats outside protected areas have been lost. Given that habitat loss and degradation are major drivers of insect declines (Forister et al. 2021; Maes & Van Dyck 2001; Sánchez-Bayo & Wyckhuys 2019; Wagner et al. 2021), it is crucial to protect and actively manage key areas. For dry prairie ecosystems, prescribed burning according to a natural fire interval (Florida FWC 2019) in a mosaic pattern that leaves unburned refugia (Schweitzer et al. 2018) is beneficial for at-risk insects. Overall, more investment is needed to conserve open-canopy prairie and savanna habitats and manage them according to natural disturbance regimes, which will help safeguard *A. loammi* and many other prairie-specialized insects.

## Supporting information

Online Resource 1

## Acknowledgements

We sincerely thank the community scientists who contributed occurrence data used in this study through iNaturalist. Without their help, this study would not have been possible. Thank you to Linda Cooper, Dean Jue, Sally Jue, Marc Minno, and Ed Perry for contributing invaluable natural history knowledge about the study species, and to the many volunteers who helped collect field data. Thank you to Dr. Vaughn Shirey and Dr. Moulay Anwar Sounny-Slitine for providing input during project conceptualization. We also thank the Geography Department at the University of Florida for allowing us to use their 32 GB RAM computer for GIS analyses. Last, we thank the Florida Museum of Natural History for supporting this project.

## Statements and Declarations

### Data availability

All code and datasets used to perform the analyses in this study, including the unobscured GPS coordinates for the *A. loammi* observations, are available on GitHub (https://github.com/Ava-C-J/LoammiCode), with the exception of the Florida Cooperative Land Cover and Florida Land Managed Areas datasets which can be located at https://myfwc.com/research/gis/wildlife/cooperative-land-cover/ and https://www.fnai.org/publications/gis-data, respectively.

