## Supplementary material for "Conservation of declining Florida ecosystems is critical for an at-risk butterfly": Online Resource 1

### Data Availability

All code and datasets used to perform the analyses in this study, including the unobscured GPS coordinates for the *A. loammi* observations, are available on GitHub (https://github.com/Ava-C-J/LoammiCode), with the exception of the Florida Cooperative Land Cover and Florida Land Managed Areas datasets which can be located at https://myfwc.com/research/gis/wildlife/cooperative-land-cover/ and https://www.fnai.org/publications/gis-data, respectively.

***Table S1*** Top 10 habitat associations of *A. loammi* ranked by habitat index when using the set of occurrence points resulting from the 150-m spatial thin (n=117). For each habitat type, index value, percent of habitat in conservation lands, and total area in acres are shown. Values for percent protected and total area are from November 2025.

| Land Cover Class | Index Value | Protected (%) | Total Area (Acres) |
|---|---|---|---|
| Dry Prairie | 17.92 | 74.77% | 152,809 |
| Palmetto Prairie | 12.94 | 66.17% | 18,798 |
| Mesic Flatwoods | 7.34 | 59.70% | 1,504,209 |
| Cultural - Riverine | 6.18 | 31.97% | 58,952 |
| Scrubby Flatwoods | 5.43 | 60.98% | 104,588 |
| Cultural - Terrestrial | 3.41 | 22.74% | 35,724 |
| Dome Swamp | 2.98 | 62.55% | 119,141 |
| Prairies and Bogs | 2.89 | 47.17% | 1,245,320 |
| Isolated Freshwater Marsh | 2.51 | 65.58% | 344,163 |
| Scrub | 2.36 | 57.53% | 163,487 |

***Table S2*** Top 10 habitat associations of *A. loammi* ranked by habitat index when using the set of occurrence points resulting from the 1-km spatial thin (n=50). For each habitat type, index value, percent of habitat in conservation lands, and total area in acres are shown. Values for percent protected and total area are from November 2025.

| Land Cover Class | Index Value | Protected (%) | Total Area (Acres) |
|---|---|---|---|
| Palmetto Prairie | 30.22 | 66.48% | 18,798 |
| Dry Prairie | 17.49 | 74.82% | 152,809 |
| Cultural - Terrestrial | 7.96 | 22.74% | 35,724 |
| Cultural - Riverine | 7.26 | 32.07% | 58,952 |
| Scrubby Flatwoods | 7.23 | 61.03% | 104,588 |
| Mesic Flatwoods | 6.40 | 59.90% | 1,504,209 |
| Mesic Hammock | 5.27 | 65.46% | 147,365 |
| Isolated Freshwater Marsh | 3.15 | 66.30% | 344,163 |
| Scrub | 2.55 | 57.63% | 163,487 |
| Prairies and Bogs | 2.34 | 48.74% | 1,245,320 |

***Table S3*** *A. loammi* occurrences (n=7) classified as > 75 m outside the top 8 natural habitat associations (excluding cultural land uses that ranked among the top 10 habitats). For each *A. loammi* occurrence, the land cover class with which it was predominately associated, the closest top 8 natural habitat, and the distance to that habitat are shown.

| Point ID | Land Cover Class | Nearest Top-8 Habitat | Distance (m) |
|---|---|---|---|
| 1 | Rural | Prairies and Bogs | 344.09 |
| 2 | Rural | Mesic Flatwoods | 330.00 |
| 3 | Tree Plantations | Mesic Flatwoods | 319.06 |
| 4 | Cultural - Terrestrial | Mesic Flatwoods | 192.35 |
| 5 | Improved Pasture | Mesic Flatwoods | 139.28 |
| 6 | Rural | Mesic Flatwoods | 80.00 |
| 7 | Improved Pasture | Isolated Freshwater Marsh | 76.16 |

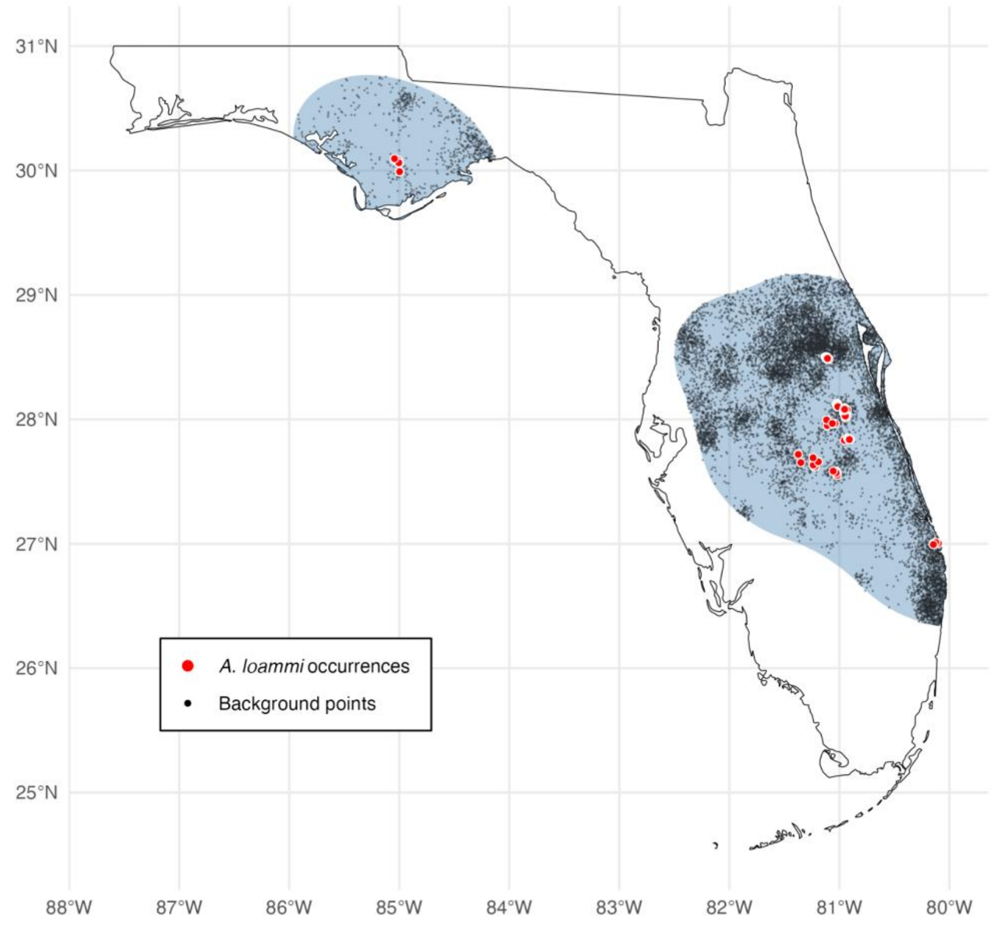


***Fig. S1*** Plot showing the accessible area estimated for *A. loammi* in blue, *A. loammi* occurrences thinned by 150 m in red (n=117), and background points weighted by butterfly sampling effort in black (n=10,000)

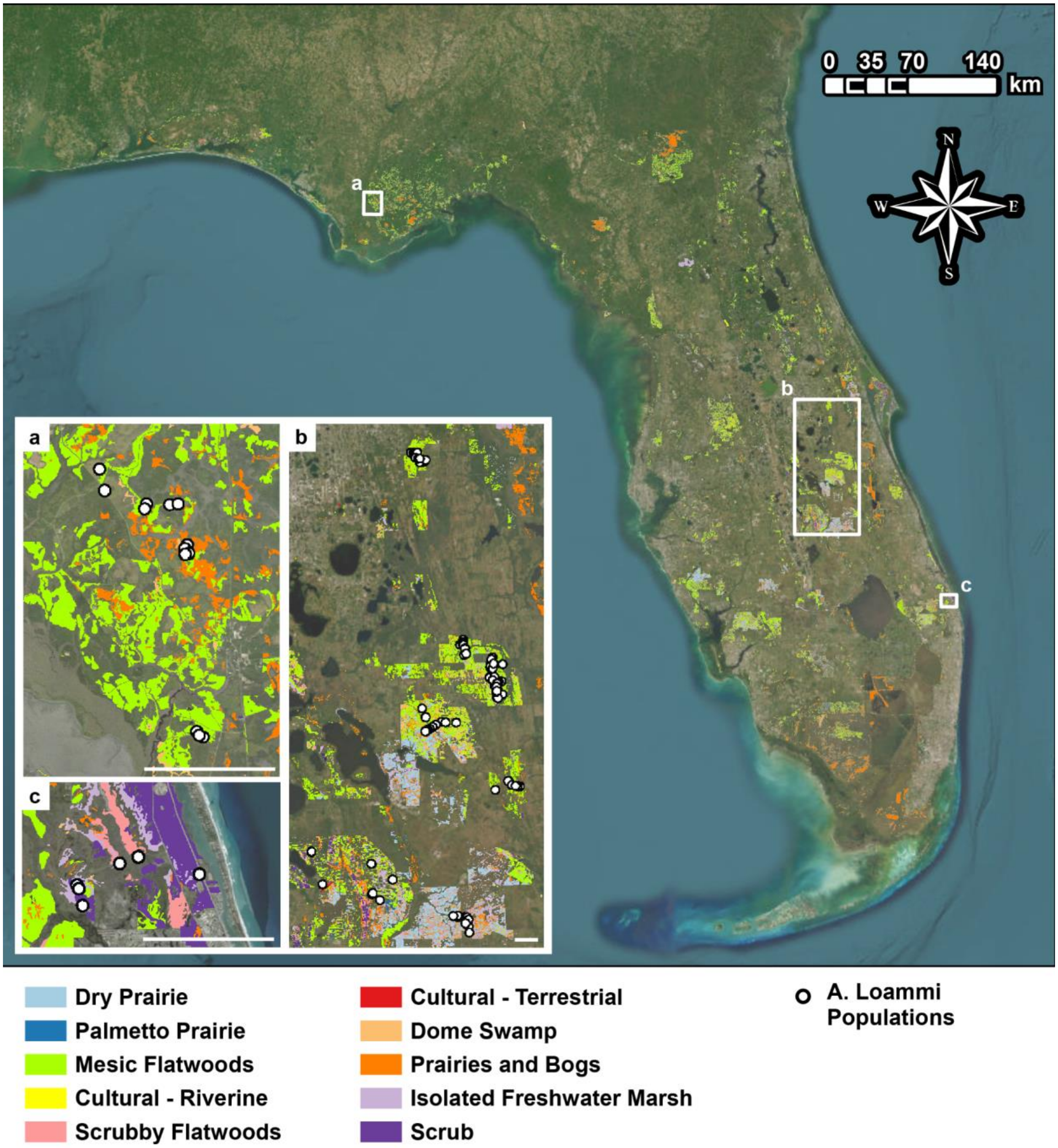


***Fig. S2*** Map of Florida showing the distributions of the top 10 *A. loammi* habitats that fall within conservation lands. Insets represent the three geographically separated Florida populations of *A. loammi*: (a) Florida panhandle, (b) central Florida, (c) south Florida. The scale bar in the lower right corner of each inset is 5 km. Maps created in ArcGIS Pro with layout completed in Adobe Illustrator. Basemap sources: Esri, Vantor, Earthstar Geographics, and the GIS User Community
